# Robust data-driven segmentation of pulsatile cerebral vessels using functional magnetic resonance imaging

**DOI:** 10.1101/2024.07.17.603932

**Authors:** Adam M. Wright, Tianyin Xu, Jacob Ingram, John Koo, Yi Zhao, Yunjie Tong, Qiuting Wen

## Abstract

Functional magnetic resonance imaging (fMRI) captures rich physiological and neuronal information that can offer insights into neurofluid dynamics, vascular health, and waste clearance function. The availability of cerebral vessel segmentation could facilitate fluid dynamics research in fMRI. However, without magnetic resonance angiography scans, cerebral vessel segmentation is challenging and time-consuming. This study leverages cardiac-induced pulsatile fMRI signal to develop a data-driven, automatic segmentation of large cerebral arteries and the superior sagittal sinus (SSS). The method was validated in a local dataset by comparing it to ground truth cerebral artery and SSS segmentations. Using the Human Connectome Project (HCP) aging dataset, the method’s reproducibility was tested on 422 participants aged 36 to 100 years, each with four repeated fMRI scans. The method demonstrated high reproducibility, with an intraclass correlation coefficient > 0.7 in both cerebral artery and SSS segmentation volumes. This study demonstrates that the large cerebral arteries and SSS can be reproducibly and automatically segmented in fMRI datasets, facilitating the investigation of fluid dynamics in these regions.

## Introduction

Functional magnetic resonance imaging (fMRI) can study various physiological and neuronal processes in the human brain [1–3]. Prior research has demonstrated the different physiological parameters influencing fMRI signals include respiration [4], cardiac pulsation [5, 6], and blood flow and blood oxygenation [7]. Specifically, the fMRI signals within cerebral vessels reflect physiological processes and potentially provide insights into cerebral vascular health [8–10] and its coupling with CSF dynamics [11]. The availability of cerebral vessel segmentation could extend fMRI research beyond gray matter to include upstream arteries and downstream veins. However, cerebral vessel segmentation is challenging and time-consuming without magnetic resonance angiography (MRA).

Time of flight (TOF) angiography is typically used to generate cerebral artery segmentations [12]. However, this imaging procedure is not always collected and is absent in many large MRI databases (e.g., Human Connectome Project, Alzheimer’s Disease Neuroimaging Initiative, UK Biobank) [13–16]. Furthermore, TOF scans are limited to arterial vessels and are not amenable to venous segmentation. Magnetic resonance venography (MRV) is used to visualize and segment venous structures and is rarely collected in MRI studies. Without angiography scans, previous studies by Tong et al. have leveraged the ratios of T1-weighted and T2-weighted anatomical scans to manually threshold the segmentations of both the internal carotid artery and superior sagittal sinus (SSS) [8, 10]. The developed techniques can individually segment cerebral arteries or veins, but no method automatically segments both vessels simultaneously. The automatic segmentation of both large cerebral arteries and veins in fMRI scans would expand the accessibility of existing MRI databases for studies that rely on segmentation of cerebrovascular regions of interest, particularly when angiography imaging is unavailable.

More importantly, even with TOF/MRV or manual segmentation on high-resolution anatomical scans, transforming the identified vessels into low-resolution fMRI space remains challenging. fMRI scans are susceptible to echo-planar imaging-induced anatomical distortions [17], which can lead to registration challenges [18]. Due to cerebral vessels’ small size, minor registration errors can significantly impact the accurate identification of vessel structures in fMRI space. Therefore, this study aimed to explore an analytical method to identify vessel regions using the fMRI data itself. By leveraging the characteristic vascular signal, we segmented vessels directly with fMRI, eliminating the need for additional scans and bypassing unnecessary procedures.

Previous research has established a consistent pulsatile signal in arterial and venous structures in cardiac-aligned fMRI. In 1999, Dagli et al. demonstrated the presence of signal fluctuations near large cerebral vessels in retrospectively aligned fMRI to the cardiac cycle [19]. Henning Voss later demonstrated that hypersampled fMRI of large cerebral vessels reveals waveforms similar to peripherally measured pulse waves [20], and Aslan et al. extend this work by extracting cardiac fMRI waveforms in the absence of finger plethysmography recordings [21]. In 2023, Hermes et al. further demonstrated the high reproducibility of cardiac-induced pulsatility in fMRI signals near large cerebral vessels, including major cerebral arteries and the SSS [22]. Thus, we were motivated to leverage this phenomenon to develop a technique that automatically segments the large cerebral arteries and SSS.

This work aims to utilize the cardiac-induced pulsatility of fMRI signals to generate a data-driven automatic segmentation of large cerebral vessels. The automatic segmentation was tested on a local dataset and the Human Connectome Project (HCP) aging cohort. In the local dataset, the automatic vessel segmentations were compared to ground truth references of the cerebral artery and SSS. Additionally, the HCP aging dataset was utilized to demonstrate the method’s robustness and reproducibility on an extensive database spanning a participant age range of 36 to 100 years. This work reveals that the large cerebral arteries and the SSS can be reproducibly and automatically segmented in fMRI datasets.

## Methods

### Human participants

Two cohorts were used to complete this study. A local cohort (n = 5, age range: 21-36 years) and an aging cohort from the Human Connectome Projects (HCP) aging 2.0 Release (n = 719, age range 36-100 years). Both cohorts contain repeated fMRI scans. All local cohort participants provided written informed consent according to procedures approved by the Institutional Committee for the Protection of Human Participants at Indiana University or Purdue University. All HCP-aging participants provided informed consent as outlined in Bookheimer et al. [13].

### Imaging acquisition

Participants underwent imaging using a 3T Prisma Siemens scanner with a 64-channel head-neck coil for the local cohort and a 32-channel head-neck coil for the HCP aging cohort. Three MR sequences were used to complete this study, including a T1-weighted (T1w) anatomical scan, a resting-state fMRI scan, and a TOF scan. The TOF imaging was only acquired for the local cohort. Simultaneous finger plethysmography (PPG) was recorded during all fMRI scans and used as a reference for retrospective cardiac alignment.

The fMRI acquisition parameters for local participants were TR = 366 msec, TE = 29.80 msec, FA = 35°, voxel size = 2.5×2.5×2.5 mm³, volumes = 500, multiband factor = 8, and an acquisition time = 3.05 minutes. All repeated local fMRI scans were acquired with anterior-posterior phase encoding. The fMRI acquisition parameters for HCP aging participants were TR = 800 msec, TE = 37.0 msec, FA = 52°, voxel size = 2.0×2.0×2.0 mm³, volumes = 488, multiband factor = 8, and an acquisition time = 6.51 minutes. The repeated fMRI scans in the HCP-aging cohort were acquired over two imaging sessions. For each session, a pair of scans with opposite phase encoding directions were acquired (anterior-posterior and posterior-anterior).

The T1w anatomical imaging data were collected using a 3D magnetization rapid gradient echo (MPRAGE) sequence. The acquisition parameters in local participants were repetition time (TR) = 2300.0 msec, echo time (TE) = 2.98 msec, flip angle (FA) = 9°, and voxel size = 1.0×1.0×1.0 mm³. The acquisition parameters for HCP aging cohorts were TR = 2500.0 msec, TE = 2.22 msec, FA = 8°, and voxel size = 0.8×0.8×0.8 mm³. The acquisition parameters for the TOF scans in local participants were TR = 21.0 msec, TE = 3.42 msec, FA = 18°, and slice thickness = 40 mm with a 2 mm gap.

### Participant image quality criteria

Each fMRI dataset was checked for finger plethysmography signal quality and motion artifacts. We developed an in-house function to automatically assess the quality of the finger plethysmography based on the percentage of signal power centered around the cardiac frequency (see Supplemental Material). fMRI data were excluded if the finger plethysmography signal was inadequate or contained motion artifact (FSL: MCFLIRT [23], maximum translation > voxel dimension). Only participants with all four fMRI repeats that passed quality checks were included in the analysis. Examples of inadequate finger plethysmography data (Supplemental Figures 1-3). No local participants were excluded from the analysis due to careful monitoring of finger plethysmography signals during data collection. We excluded 297 HCP participants from the analysis (101 had at least one of the scans without a plethysmography recording, 13 were excluded for both plethysmography and motion, 19 were excluded for excessive motion, and 164 were excluded for poor plethysmography data). Following data quality control, five local participants and 422 HCP aging participants were included in the analysis.

### Image processing

#### fMRI pre-processing

The initial ten volumes of the fMRI were excluded to reduce T1-relaxation effects. A voxel-wise high-pass Butterworth filter with a cutoff frequency of 0.005 Hz was then applied to remove the fMRI signal’s DC offset. Although flipped phase-encoding data was available, distortion correction was not performed because it could disrupt the slice timing profile due to data interpolations between adjacent slices with different readout times.

#### General vessel region identification

An initial overly inclusive cerebral artery and SSS region were generated to define a general search region in the data-driven vessel segmentation. These search regions are referred to as the general cerebral artery region of interest (ROI) and general SSS ROI. The general cerebral artery ROI was derived from the statistical brain atlas Dunås et al. developed using manually labeled 4D flow MRI [24]. The atlas was modified to include the basilar artery, posterior cerebral artery (PCA), anterior cerebral artery (ACA), middle cerebral artery (MCA), and their major branches. The mask was binarized in MNI space by selecting all voxels with a probability greater than 0.5% to create an overly inclusive ROI around the major cerebral arteries. A general SSS ROI was generated by manually segmenting the SSS in the MNI152 T1 1×1×1 mm^3^ brain, which was later dilated in the subject’s fMRI space to make it overly inclusive. Subsequently, the statistical brain atlas and manual SSS segmentation were transformed from the MNI space to the T1w space (ANTS: antsApplyTransforms, non-linear interpolation) and then to the fMRI space (FSL: FLIRT, nearest-neighbor interpolation). Following the SSS mask transformation into fMRI space, it was dilated (Matlab: imdilate, kernel size = 3) to ensure the inclusiveness of the general SSS ROI. Representative examples of the general arterial and SSS ROIs are displayed in the lower left panels of Figure 1, 3.B.

**Figure 1.**
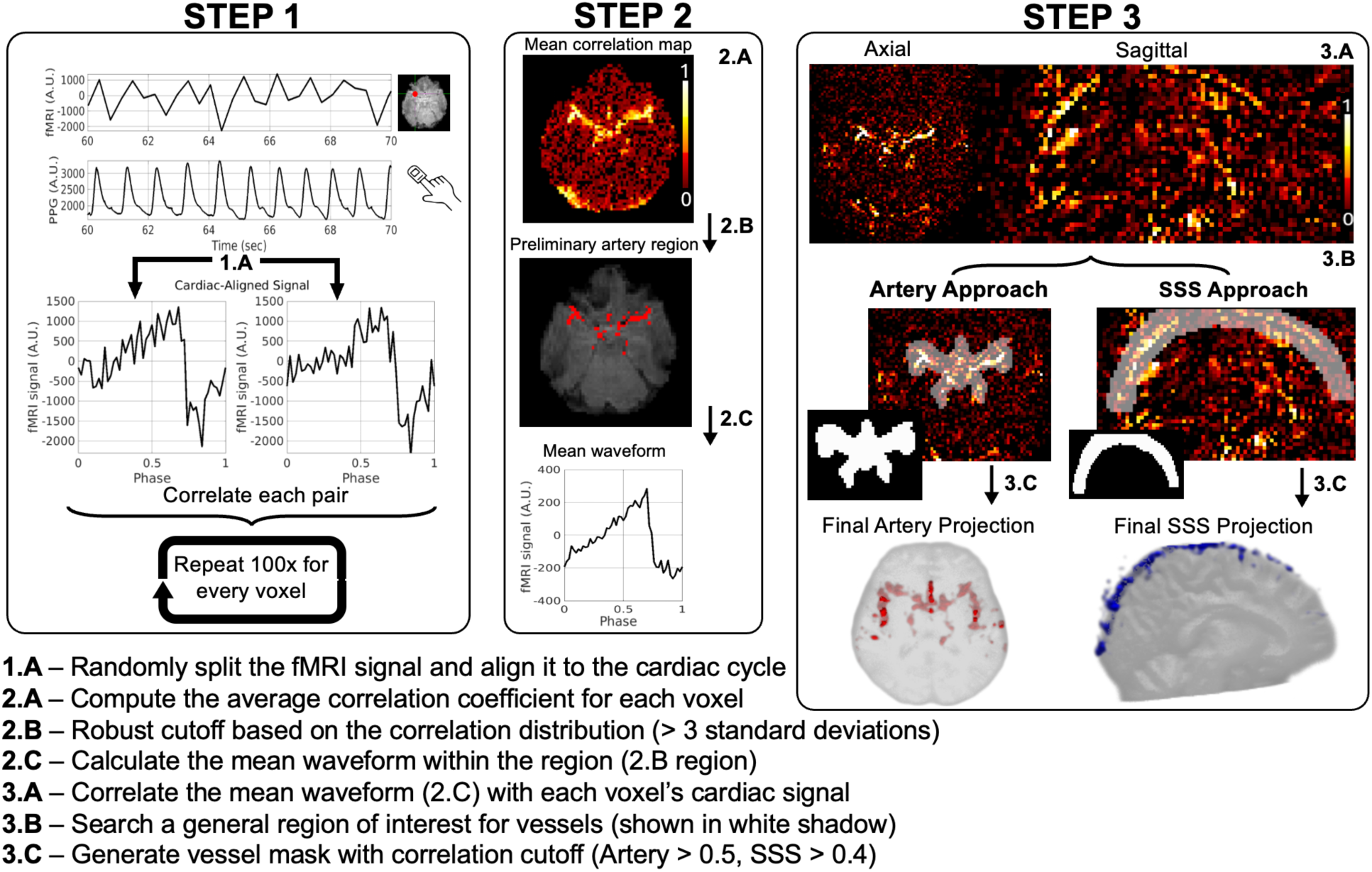
Schematic overview of the data-driven segmentation pipeline. (1.A) Time-detrended single voxel fMRI time series (top), the corresponding finger plethysmography signal (middle), and cardiac-aligned signal pair from an arterial voxel (bottom). (2.A) Mean correlation map computed from the 100 signal pair correlations completed in Step 1. (2.B) Preliminary artery region generated by identifying voxels with correlations three standard deviations above the mean. (2.C) The mean waveform is based on the voxels in the preliminary artery region. (3.A) Correlation map between the mean waveform of the preliminary artery region and the voxel-wise cardiac-aligned fMRI signal. (3.B left) General artery ROI overlay with the correlation map (3.B right) and general SSS ROI overlay with the correlation map. (3.C left) The final artery segmentation (displayed as a 3D projection) is defined as voxels within the general artery ROI and correlations greater than 0.5. (3.C right) The final SSS segmentation (displayed as a 3D projection) is refined using an SSS-specific iteration approach detailed in the methods. **Abbreviations:** fMRI – functional magnetic resonance imaging, SSS – superior sagittal sinus, ROI – region of interest.

### Data-driven vessel segmentation generation

The data-driven vessel segmentation is a multi-step process that includes three significant steps. These steps are summarized in Figure 1 and include a randomized realignment of the fMRI data to the cardiac cycle (Step 1), the generation of a preliminary artery region (Step 2), and the generation of a finalized segmentation for both cerebral arteries and the SSS (Step 3).

#### Randomized fMRI signal cardiac alignment (Step 1)

The preprocessed fMRI signals were randomly split into two temporal subsets and retrospectively aligned to the cardiac cycle, resulting in a voxel-wise cardiac-aligned signal pair (Figure 1, 1A). The splitting was completed by randomly selecting half of the time points of the time series using (Matlab: randperm) to represent the first temporal set. The second temporal set comprised all time points not used in the first. After selecting the time points, each temporal set was realigned to the cardiac cycle using finger plethysmography as the reference. Then, a voxel-wise correlation coefficient between each cardiac-aligned signal pair was calculated. The process of generating cardiac-aligned signal pairs and their correlation coefficient was repeated one hundred times.

#### Preliminary artery region (Step 2)

A voxel-wise mean correlation map was generated by taking the mean of the one hundred repeated correlation coefficients (Figure 1, 2A). A higher correlation coefficient indicates an increased likelihood that the voxel contains a vessel due to the strong pulsatility [22]. Considering the correlation coefficients vary among participants, a data-driven approach was utilized to identify a participant-specific correlation threshold to define highly likely vascular voxels. The threshold was generated as the correlation coefficient three standard deviations above the mean of all brain voxels (T_preliminary_). Once this threshold was generated, all voxels within the general cerebral artery ROI with mean correlation coefficients greater than T_preliminary_ were considered probable vascular voxels. This process produced the preliminary artery region that will be later refined (Figure 1, 2B). The preliminary artery region represents highly pulsatile cardiac-aligned fMRI signals, mainly consisting of actual vessels with some spurious voxels. The mean waveform of this preliminary artery region was then calculated to represent the signal waveform of the arterial region (Figure 1, 2C). This waveform was used to refine future cerebral artery and SSS segmentations.

#### Refined vessel segmentation (Step 3)

In the final step, a new correlation map was generated by calculating a voxel-wise correlation between the voxel-wise cardiac-aligned fMRI signal and the mean waveform from the preliminary artery region (Figure 1, 3A). This correlation map was thresholded to generate final segmentations of the cerebral arteries and SSS but with slightly different approaches. In the artery segmentation approach, voxels within the general artery ROI (Figure 1, 3B left) with correlation coefficients greater than 0.5 were selected for the final artery segmentation (Figure 1, 3C left).

We observed that the SSS pulsates at a slight time delay compared to arteries, and the SSS mask was refined with the following steps. In step-SSS_1, voxels within the general SSS ROI (Figure 1, 3B right) and with a higher correlation coefficient (greater than 0.4) were selected as probable SSS voxels. Afterward, in step-SSS_2, an iterative approach was used to finalize the SSS segmentation. In this iterative approach, the mean waveform of probable SSS voxels was calculated and used to generate an SSS correlation map by correlating with the voxel’s cardiac-aligned fMRI signal. Step-SSS_1 (using the most recent probable SSS voxels as the starting segmentation) and step-SSS_2 were repeated until the new SSS segmentation volume was stabilized, defined by a volume percent change of less than 10% compared to the previous iteration, resulting in the final SSS segmentation (Figure 1, 3C right).

### Performance assessment and reproducibility

#### Qualitative comparison with ground truth

Ground truth artery regions were generated using TOF scans. Manually set intensity thresholds were used to binarize TOF scans, which were then quality-checked to ensure proper segmentations of the PCA, ACA, and MCA and their major branches. The binarized TOF scans were transformed into T1w space (FSL: FLIRT, nearest-neighbor interpolation) and then to fMRI space (FSL: FLIRT, nearest-neighbor interpolation). Once in fMRI space, the TOF binary mask was limited to voxels within the general cerebral artery ROI (to ensure the overall segmentation region matched the one used in the data-driven segmentation) and was defined as the arterial ground truth segmentation. Ground truth SSS were manually drawn on T1w images and were transformed into fMRI space (FSL: FLIRT, nearest-neighbor interpolation). Once in fMRI space, the manually drawn SSS segmentation was limited to voxels within the general SSS ROI (to ensure the overall segmentation region matched the one used in the data-driven segmentation) and was defined as the SSS ground truth segmentation.

The data-driven artery segmentation results of the local participants were qualitatively compared with the corresponding ground truth artery segmentation for the five local participants (the HCP-aging dataset does not have TOF scans). The SSS segmentation results of both local and HCP participants were qualitatively compared with the corresponding ground truth SSS segmentation (Local: n = 5, HCP, n = 5).

#### Reproducibility assessment

The reproducibility of the data-driven segmentation was assessed by calculating the intraclass coefficient (ICC, two-way mixed effects, single rater, absolute agreement) of the segmentation volumes. All participants with four scans that passed the quality assessment step were used in this calculation (Local: n = 5, HCP, n = 422).

## Results

### Data-driven and ground truth segmentation results align

The ground truth (left of image pair) and data-driven (right of image pair) segmentations of five local and five HCP-aging participants are summarized in Figure 2. All segmentations were displayed as projections and overlaid onto T1-weighted images in fMRI space (arteries – axial view, SSS – sagittal view). Qualitatively, the data-driven artery segmentation matches closely with the ground truth segmentations derived from TOF images (Figure 2 – top). In all participants, the Circle of Willis and its components are distinguishable and comparable to their TOF comparisons. In the fourth local participant, the right side of the arterial data-driven segmentation had fewer segmented voxels. The reason behind this could not be determined. In the local (Figure 2 – middle) and HCP (Figure 2 – bottom) participants, the SSS segmentation results show a continuous vessel structure from the anterior to the posterior in the 3D projection and closely align with the manually segmented ground truth.

**Figure 2.**
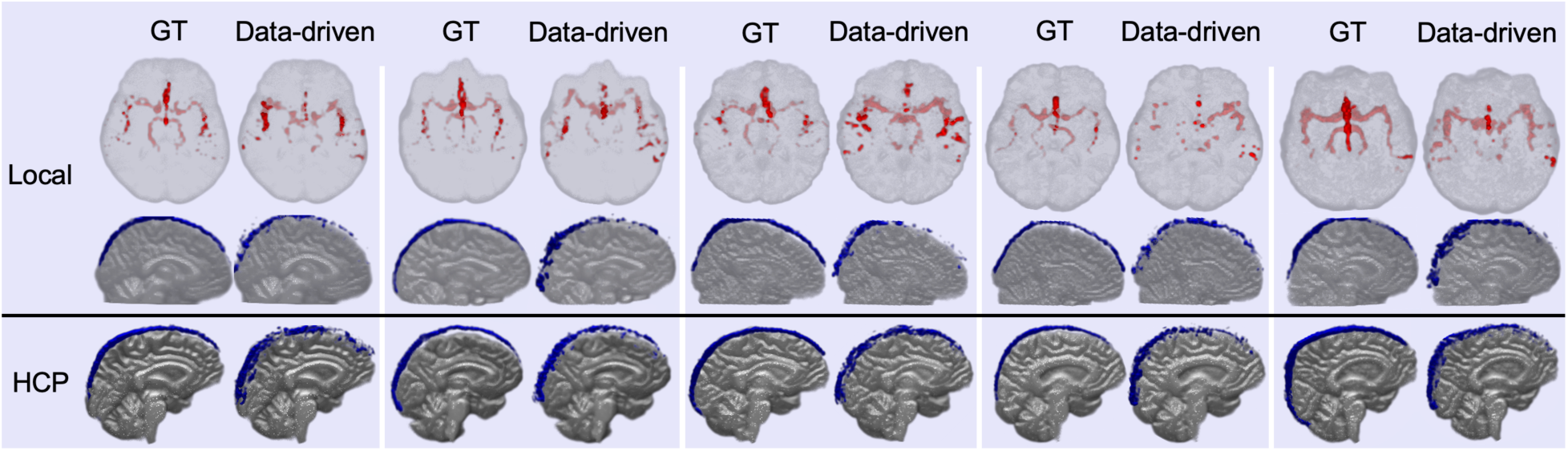
3D projections of fMRI-derived vessel segmentations (Data-driven, right of pair) in comparison with ground truth (GT, left of pair) segmentations in fMRI space. For local participants, the data-driven arterial segmentations are compared with TOF intensity-based arterial segmentations (top row), and the data-driven SSS segmentations are compared with manual SSS segmentations (middle row). For five representative HCP participants, the data-driven SSS segmentations are compared with manual SSS segmentations (bottom row). **Abbreviations:** fMRI – functional magnetic resonance imaging, GT – ground truth, TOF – time of flight, SSS – superior sagittal sinus, HCP – Human Connectome Project.

### Data-driven segmentations are reproducible

The data-driven segmentation of repeated scans for a local participant (age 35), a younger HCP participant (age 40), and an older HCP participant (age 74) are displayed in Figure 3. For each repeated scan, one axial slice of the artery segmentation (red) is overlaid onto T1-weighted MR images (Figure 3A), and one sagittal slice of the SSS segmentation (blue) is overlaid onto T1-weighted MR images (Figure 3B). The qualitative comparison of the four repeated scans demonstrates that the data-driven arterial and SSS segmentations perform consistently across all repeats. The single-slice views display some discontinuity in the segmentation that is not visible when viewing the segmentations as a projection.

**Figure 3.**
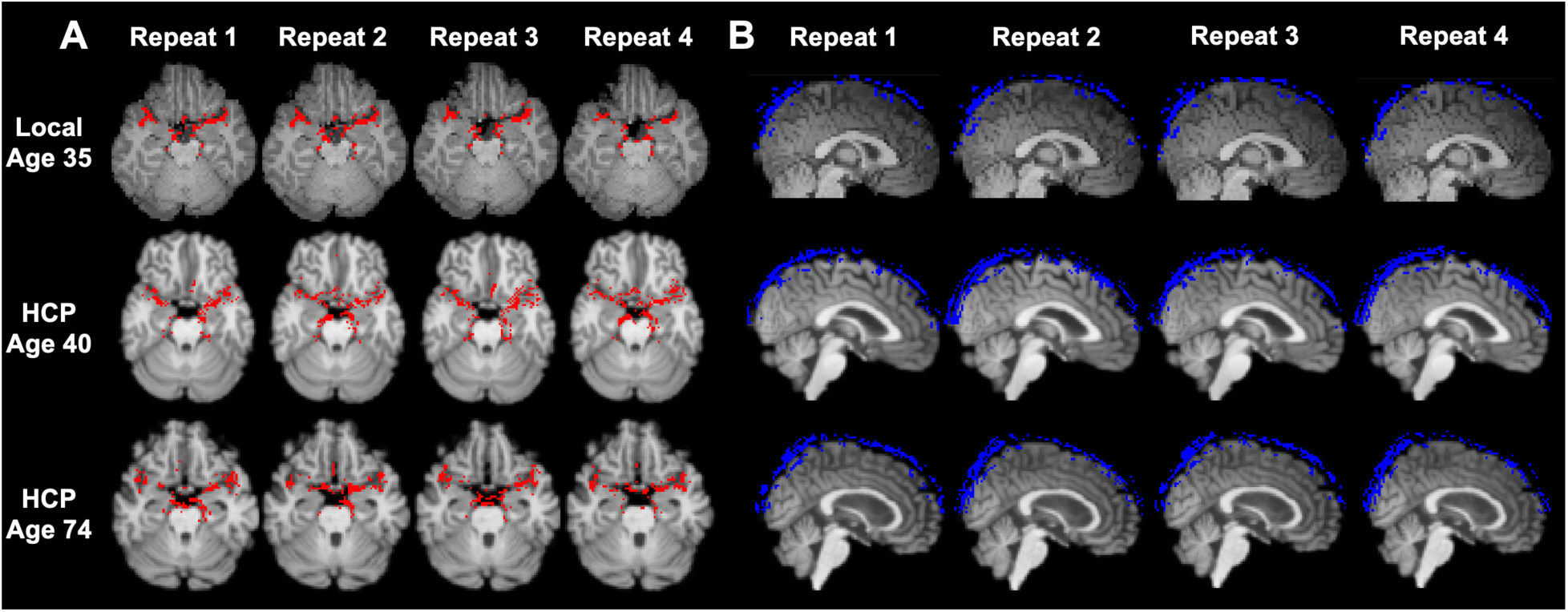
Representative data-driven arterial and SSS segmentations of repeated scans. The three representative participants included were a local participant (35 years, top), one younger HCP participant (40 years, middle), and one older HCP participant (74 years, bottom). (A) A single axial slice artery segmentation (red) includes portions of the middle cerebral and posterior cerebral arteries. (B) A single sagittal slice SSS segmentation (blue). **Abbreviations:** SSS – superior sagittal sinus, HCP – Human Connectome Project.

The comparison of the segmentation volume between repeats is illustrated in Figure 4 (A – Arterial, B – SSS). The repeated arterial segmentation volumes had an ICC of 0.798 (95% CI: 0.755 – 0.834, p<0.001). The repeated arterial segmentation volumes had an ICC of 0.734 (95% CI: 0.698 – 0.768, p<0.001). The ICC above 0.7 in both segmentation approaches demonstrates good reproducibility across scans.

**Figure 4.**
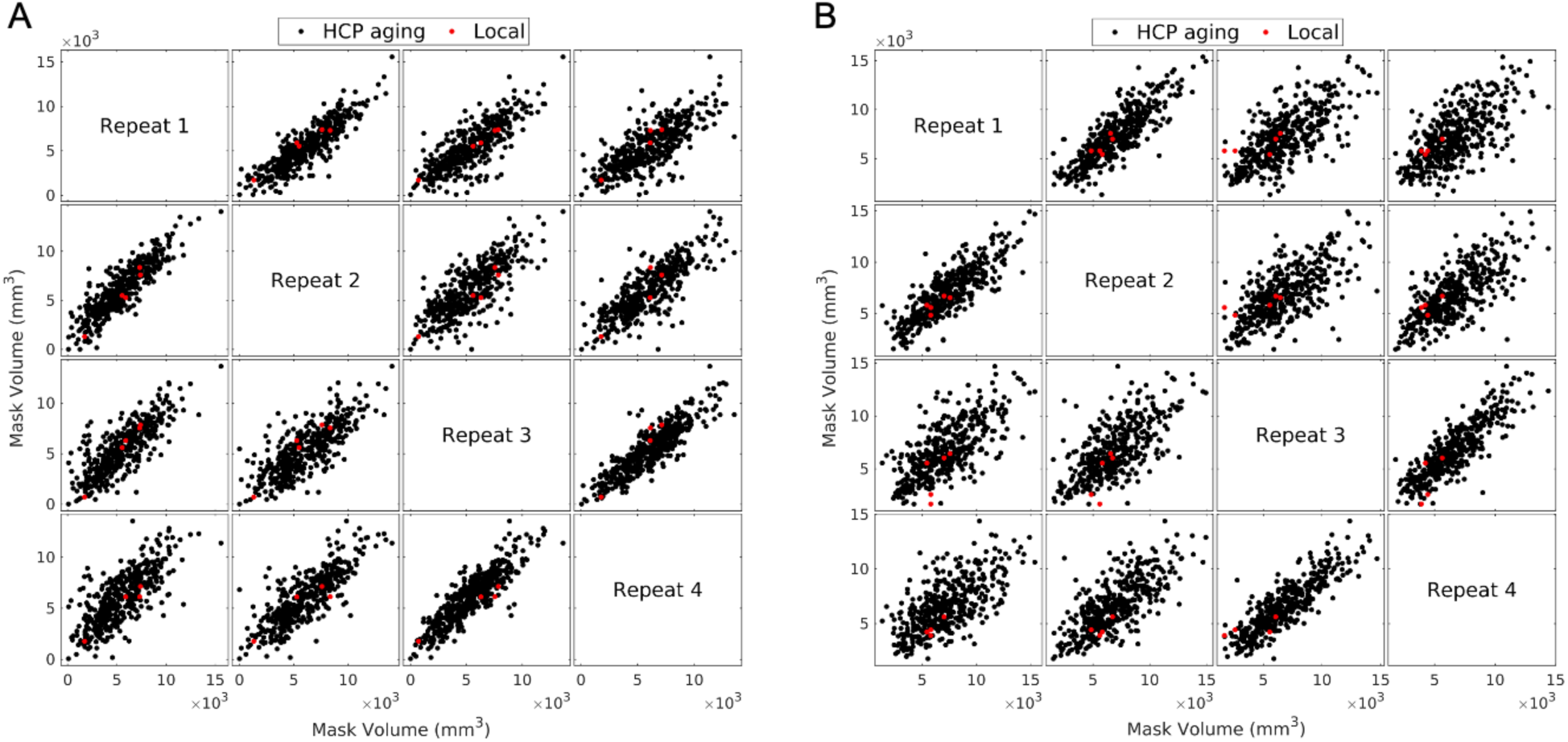
Gridded scatter plots of the mask volumes compared to respective repeated scans. (A) Comparison of repeated arterial segmentation volumes for local (red) and HCP (black) participants. (B) Comparison of repeated SSS segmentation volumes for local (red) and HCP (black) participants. **Abbreviations:** HCP – Human Connectome Project, SSS – superior sagittal sinus.

**Figure 5.**
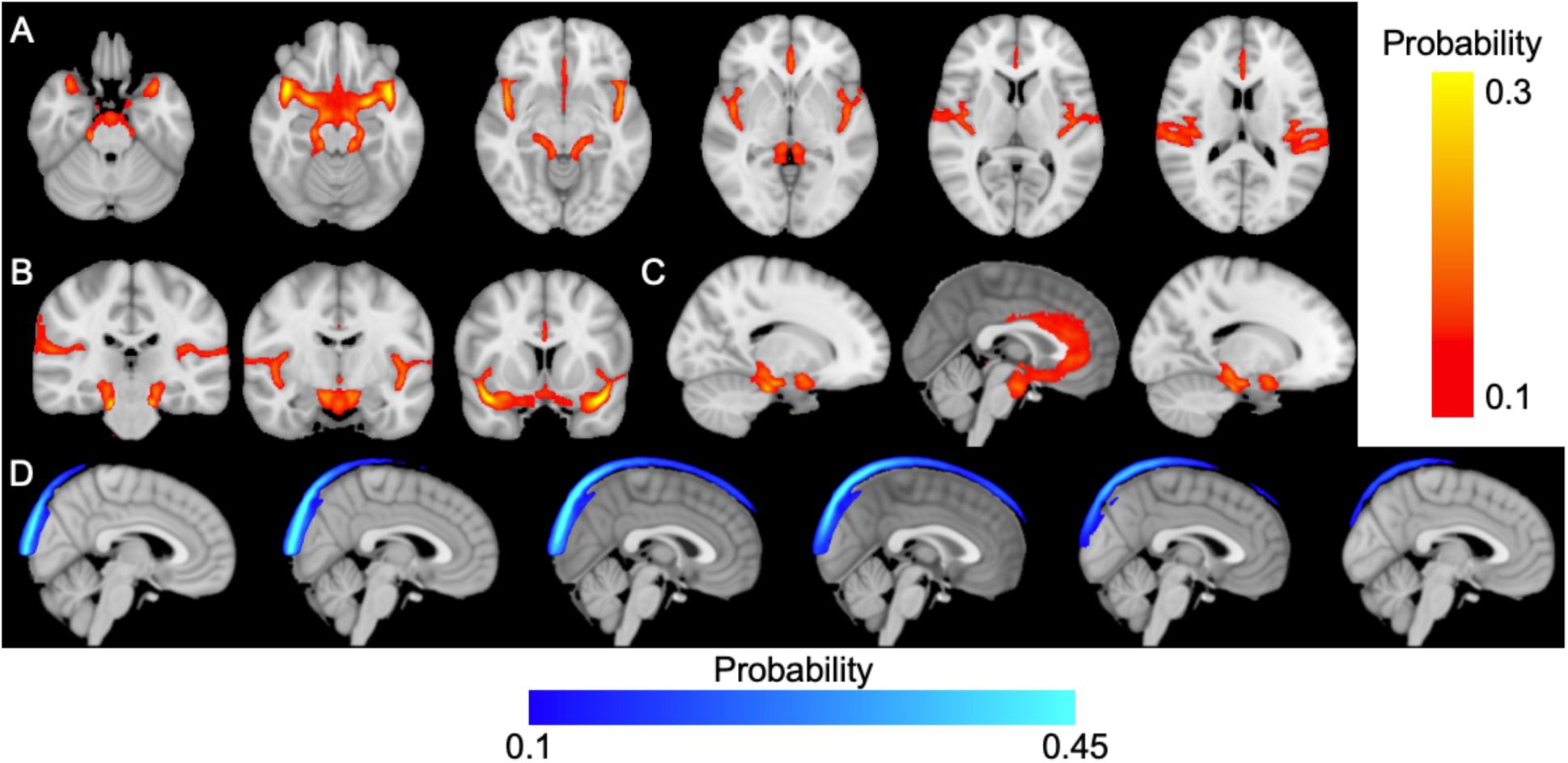
HCP participants’ average probability of large cerebral arteries. (A-C) and superior sagittal sinus (D) segmentations in MNI152 T1 1 mm isotropic space. **Abbreviations:** HCP – Human Connectome Project.

### Group averaged cerebral artery and SSS segmentations in MNI space

The large cerebral artery and SSS segmentations for all HCP participants’ scans were transformed into the MNI152 T1 1 mm space and averaged to create a group probability map of the arteries (Figure 6A-C) and SSS (Figure 6D). The highest observed probability in the arterial space was approximately 30% and observed near the ascending portion of MCA-m1 before the MCA bifurcation (Figure 6B – right panel). The MCA-m1 segment had higher probabilities than the segments of the ACA and PCA. The highest probability in the SSS space was approximately 50% and observed near the posterior portion of the SSS (Figure 6D).

**Figure 6.**
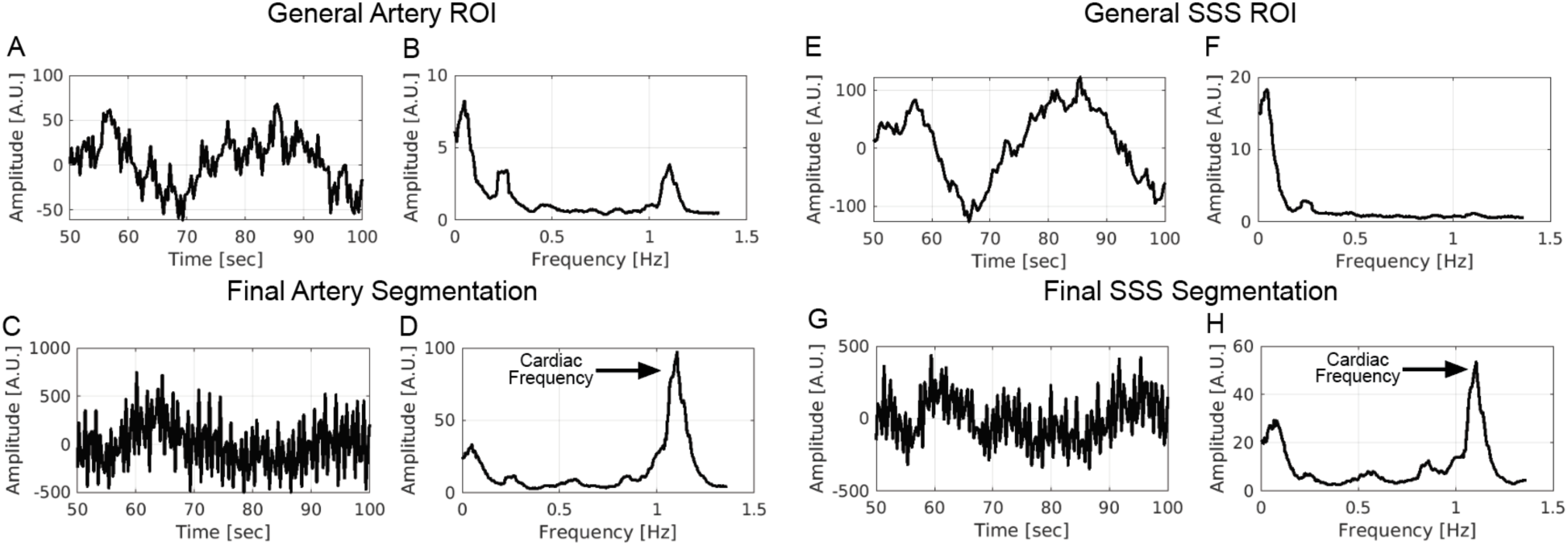
A representative participant’s progression of the mean fMRI time series and frequency response from the start of the segmentation process using the general vascular ROI to the end of the data-driven segmentation (final vascular segmentation). In this scan, the fMRI repetition time was 0.366 seconds (Nyquist frequency = 1.366 Hz), and the participant’s average heart rate was 67 beats per minute (cardiac frequency = 1.117 Hz). The general artery ROI mean time series (A) and frequency spectrum (B). The final artery segmentation mean time series (C) and frequency spectrum (D). The general SSS ROI mean time series (E) and frequency spectrum (F). The final SSS segmentation mean time series (G) and frequency spectrum (H). **Abbreviations:** fMRI – functional magnetic resonance imaging, ROI – region of interest, SSS – superior sagittal sinus.

### Spectral analysis confirms that data-driven segmentation pinpoints voxels with high cardiac pulsatility

Data-driven segmentation is a multiple-step process that starts with a general vessel ROI and is finalized with vascular segmentation. In a representative subject, for the general vascular ROI and the final vascular segmentation, the mean fMRI real-time signal and the corresponding frequency response are summarized in Figure 6. The general artery ROI time series contains some high-frequency fluctuations (Figure 6A), but the magnitude of this fluctuation is heavily amplified in the final artery segmentation (Figure 6C). This observation is further supported by the increase in the amplitude of the cardiac frequency (∼1 Hz) seen from the general artery ROI (Figure 6B) to the final artery segmentation (Figure 6D). These findings are mirrored in the SSS segmentation results. The general SSS ROI time series contains little high-frequency fluctuations (Figure 6EF), while the final SSS segmentation contains high cardiac frequency activity (Figure 6GH). These observations confirm that the data-driven segmentation effectively pinpoints voxels with high cardiac pulsatility in both the arteries and SSS.

## Discussion

This work utilized the pulsatile fMRI signal within large vessels to automatically segment major cerebral arteries and the SSS. The comparison to TOF segmentations confirmed the accuracy of the data-driven fMRI method in identifying large cerebral arteries. The implementation of this technique within the HCP-aging dataset further demonstrated its high reproducibility in same-day, between-day, and opposite-phase encoding scans. Additionally, spectral analysis confirmed that the algorithm identified voxels with highly pulsatile signals at the cardiac frequency in both arterial and venous regions. The ability to automatically segment large cerebral vessels facilitates future investigations of hemodynamics in these regions.

Physiological processes, such as cardiac pulsation and respiration, are well documented to impact fMRI signals [4, 5, 25–27]. The primary contrast in fMRI, the Blood Oxygen Level Dependent (BOLD) signal, is related to blood [2]. These physiological processes affect cerebrovascular dynamics more directly than neuronal activation. Therefore, studying cerebrovascular dynamics using fMRI is natural and sometimes advantageous. Recently, fMRI has been used to investigate phenomena such as blood flow and blood volume [7], cerebral transit time [8, 10, 28], and cerebral vascular reactivity [29–32]. Extracting the time series from large vessels in fMRI data is the first step in many of these studies. The ability to automatically identify large vessels in fMRI could significantly benefit studies focused on cerebral hemodynamics that lack TOF imaging. More importantly, it avoids registration errors and alleviates the time constraints of manually segmenting vascular regions.

In addition to promoting the investigation of vascular hemodynamics, identifying vessel regions will also benefit traditional fMRI analysis. The influence of vascular signal is considered a “noise source” in conventional fMRI analysis [3, 33], where the primary focus is on neuronal activation. In fMRI, signals from arteries and veins are seen as non-neuronal physiological signals, which can interfere with regional neuronal activation (i.e., from neurovascular coupling) through the cerebrovasculature. The noise has been demonstrated to be highly present in vasculature at the following frequencies, including the low-frequency (0.01-0.1 Hz) [6, 34], respiratory (0.2-0.3 Hz) [3, 35, 36], and cardiac (0.9-1.5 Hz) frequencies [19, 35, 37]. Moreover, Zhong et al. demonstrated that large arteries and veins significantly impact the surrounding voxels due to the blood’s magnetic properties, thus confounding the functional connectivity measures [38]. This underscores the necessity of excluding large vessel signals to complete reliable connectivity analyses [38]. Automatic segmentation of large vessels in fMRI space can enhance denoising techniques in two ways: 1) by accurately extracting physiological noise by regressing out large vessel temporal series, and 2) by avoiding regions with high vascular density in connectivity analysis.

This technique holds certain limitations warranting future exploration. One limitation is that the technique requires simultaneous finger plethysmography. While finger plethysmography is an easy signal to acquire, it is not always available in all datasets. In future work, techniques can be implemented to generate finger plethysmography signals using fMRI data [21, 39] and complete these segmentations without simultaneous finger plethysmography recordings. Another limitation is that the segmentation relies on signals induced by cardiac pulsation, and it is unclear how it will perform in vascular pathologies such as stroke. Lastly, this technique identifies vascular regions in fMRI space, and due to fMRI’s inherently large voxel size, the segmentations may contain partial volumes of cerebrospinal fluid, gray matter, and white matter.

In conclusion, this work presents a robust, automatic, data-driven tool to segment arteries and veins in fMRI datasets. This approach provides a valuable tool for large vessel segmentations in fMRI, which can be completed independent of vascular angiography scans and, therefore, without registration errors. By offering a reliable method to discern vascular components, the tool is a crucial resource for improving the accuracy of denoising models used in conventional fMRI connectivity analyses and enhances the feasibility of conducting detailed hemodynamic studies.

## Funding

This work was supported by the National Institutes of Health grants: F30AG084336 (PI: Adam Wright) and RF1AG083762 (PI: Qiuting Wen). Data acquisition was supported in part by NIH grant S10OD012336. Additionally, research reported in this publication was supported by the National Institute on Aging of the National Institutes of Health under Award Number U01AG052564 and by funds provided by the McDonnell Center for Systems Neuroscience at Washington University in St. Louis.

## Code and data availability

The code to complete the data-driven segmentation is available upon request and will be available on a GitHub repository following peer-reviewed publication.

Some of this study’s data came from the HCP-Aging 2.0 Release data, DOI: 10.15154/1520707.

## Authors’ Contributions

AMW, TX, QW, and YT conceptualized the study. AMW, TX, QW, YT, and JI drafted the article. AMW, TX, QW, YT, JI, JK, and YZ revised the article. AMW and QW collected local datasets. AMW, JK, and YZ performed statistical analyses. AMW, TX, QW, YT, JI, JK, and YZ interpreted the results.

## Competing Interests

The author’s do not have any competing interests to disclose.

## Supporting information

see Supplemental Material

## Notes

https://www.humanconnectome.org/study/hcp-lifespan-aging/document/hcp-aging-20-release

### Competing Interest Statement

The authors have declared no competing interest.

