## Supplementary material for "Robust data-driven segmentation of pulsatile cerebral vessels using functional magnetic resonance imaging": see Supplemental Material

### Finger plethysmograph data quality check.

An in-house algorithm implementing a power spectrum analysis was used to determine the quality of the finger plethysmography data (Supplemental Figures 1-3). The algorithm automatically detected the cardiac frequency and calculated the percentage of power within $\pm 0.15$ Hz of the cardiac frequency compared to the total power from -0.15 Hz of the cardiac frequency to -0.15 Hz of the first cardiac frequency harmonic (Equation 1).

$$Plethsmography\,\% = \frac{Power\ within\ \pm 0.15\ Hz\ of\ the\ Cardiac\ Freq.}{Power\ within - 0.15\ of\ the\ Cardiac\ Freq.\ to\ -0.15\ Hz\ of\ the\ first\ Cardiac\ Harmonic\ Freq.} \quad (1)$$

It was empirically determined that a cutoff of 80% was sufficient to remove poor finger plethysmography data. Considering every scan in the HCP aging dataset with a plethysmography file available (number of scans = 2661), this cutoff removed 456 scans (~17%). This is comparable to another study using HCP data, which determined ~12% of finger plethysmography data was unusable [1].

### Example of useable finger plethysmography with high data quality.

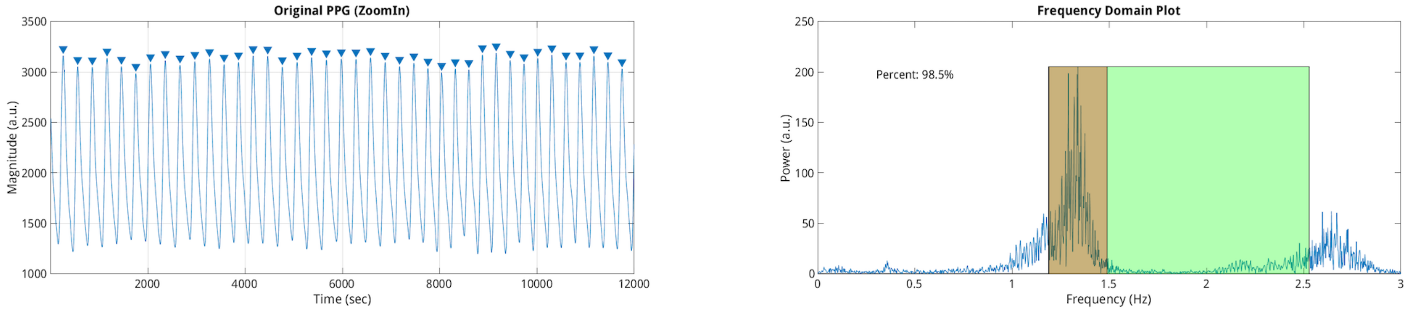


**Supplemental Figure 1:** Usable finger plethysmography with a zoomed-in peak detection of heartbeats (left). The power spectrum of the plethysmography curve where the plethysmography percentage using equation 1 was 98.5% (right).

### Example of useable finger plethysmography with intermediate data quality.

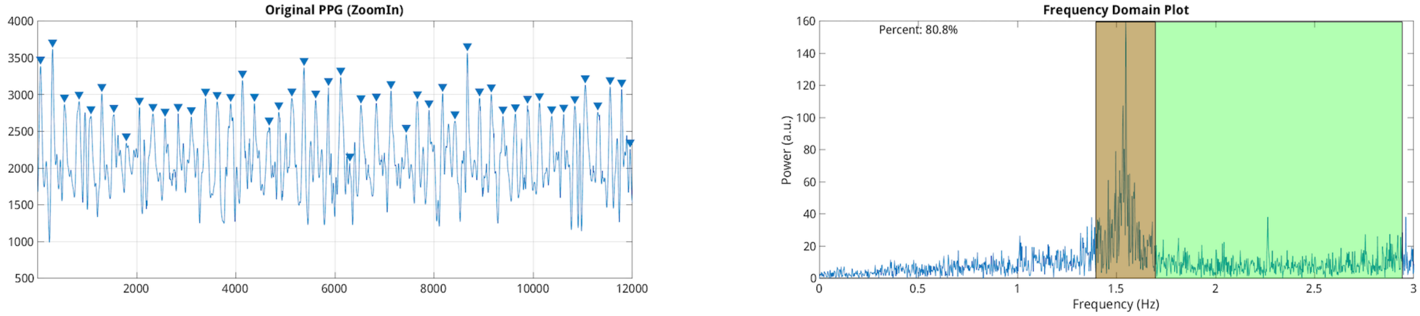


**Supplemental Figure 2:** Usable finger plethysmography with a zoomed-in with some potential for errors in peak detection of heartbeats (left). The power spectrum of the plethysmography curve where the plethysmography percentage using equation 1 was 80.8% (right)

### Example of unusable finger plethysmography with poor data quality.

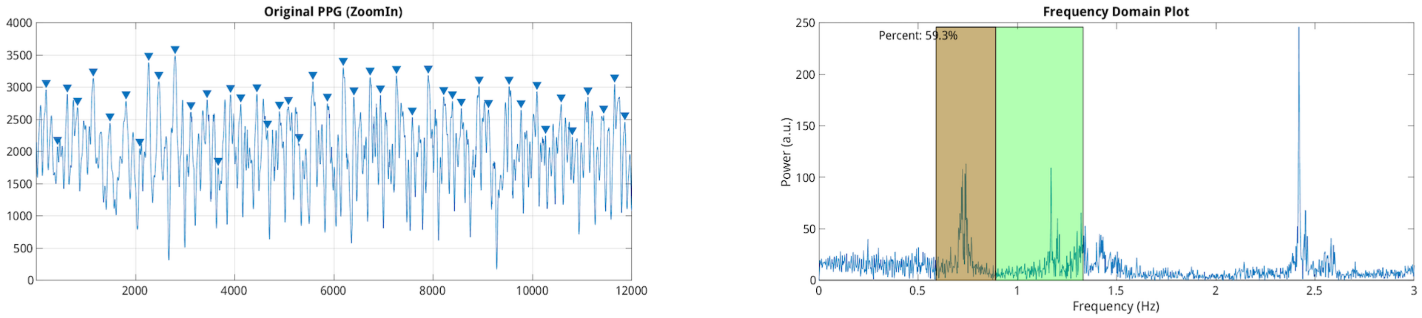


**Supplemental Figure 3:** Unsable finger plethysmography with a zoomed-in view of inaccurate peak detection of the heartbeats(left). The power spectrum of the plethysmography curve where the plethysmography percentage using equation 1 was 59.3% (right)
